# A high-quality pseudo-phased genome for *Melaleuca quinquenervia* shows allelic diversity of NLR-type resistance genes

**DOI:** 10.1101/2023.04.27.538497

**Authors:** Stephanie H Chen, Alyssa M Martino, Zhenyan Luo, Benjamin Schwessinger, Ashley Jones, Tamene Tolessa, Jason G Bragg, Peri A Tobias, Richard J Edwards

## Abstract

**Background:** The coastal wetland tree species *Melaleuca quinquenervia* (Cav.) S.T.Blake (Myrtaceae), commonly named the broad-leaved paperbark, is a foundation species in eastern Australia, Indonesia, Papua New Guinea, and New Caledonia. The species has been widely grown as an ornamental, becoming invasive in areas such as Florida in the United States. Long-lived trees must respond to a wide range pests and pathogens throughout their lifespan, and immune receptors encoded by the nucleotide- binding domain and leucine-rich repeat containing (NLR) gene family play a key role in plant stress responses. Expansion of this gene family is driven largely by tandem duplication, resulting in a clustering arrangement on chromosomes. Due to this clustering and their highly repetitive domain structure, comprehensive annotation of NLR encoding genes within genomes has been difficult. Additionally, as many genomes are still presented in their haploid, collapsed state, the full allelic diversity of the NLR gene family has not been widely published for outcrossing tree species.

**Results:** We assembled a chromosome-level pseudo-phased genome for *M*. *quinquenervia* and describe the full allelic diversity of plant NLRs using the novel FindPlantNLRs pipeline. Analysis reveals variation in the number of NLR genes on each haplotype, differences in clusters and in the types and numbers of novel integrated domains.

**Conclusions:** We anticipate that the high quality of the genome for *M. quinquenervia* will provide a new framework for functional and evolutionary studies into this important tree species. Our results indicate a likely role for maintenance of NLR allelic diversity to enable response to environmental stress, and we suggest that this allelic diversity may be even more important for long-lived plants.

## Background

*Melaleuca quinquenervia* (Cav.) S.T. Blake is a broad-leaved paperbark tree endemic to the wetlands of eastern Australia, Papua New Guinea, New Caledonia and Indonesia (Figure 1) [1]. *M. quinquenervia* belongs to the family Myrtaceae, a large family of woody flowering plants consisting of over 144 genera and 5,500 species [2] with the genus *Melaleuca* comprising almost 300 species [1]. While *M. quinquenervia* is keystone species in its endemic range, it is also planted extensively as an ornamental tree and is one of the main commercial species of the genus *Melaleuca*, as a source of essential oils and nectar for honey [1]. *M. quinquenervia* is highly invasive in the wetlands of Florida in the United States after the arrival of the species as an ornamental plant in the early 1900s [3]. Since its introduction, it has caused significant loss of native vegetation and associated biodiversity as well as increased fire risk in wetland areas [4]. The management of *M. quinquenervia* outside its endemic range has a serious economic impact due to labour intensive management practices including site monitoring, the physical removal of trees, and herbicide application [3]. High accuracy reference genomes are important for molecular and evolutionary studies, as well as providing a tool for strategic management of native and invasive species. With no current genome resource for *M. quinquenervia*, molecular research has been limited to homology-based studies using plants within the Myrtaceae family, including the closely-related species *Melaleuca alternifolia* [5–7].

**Figure 1.**
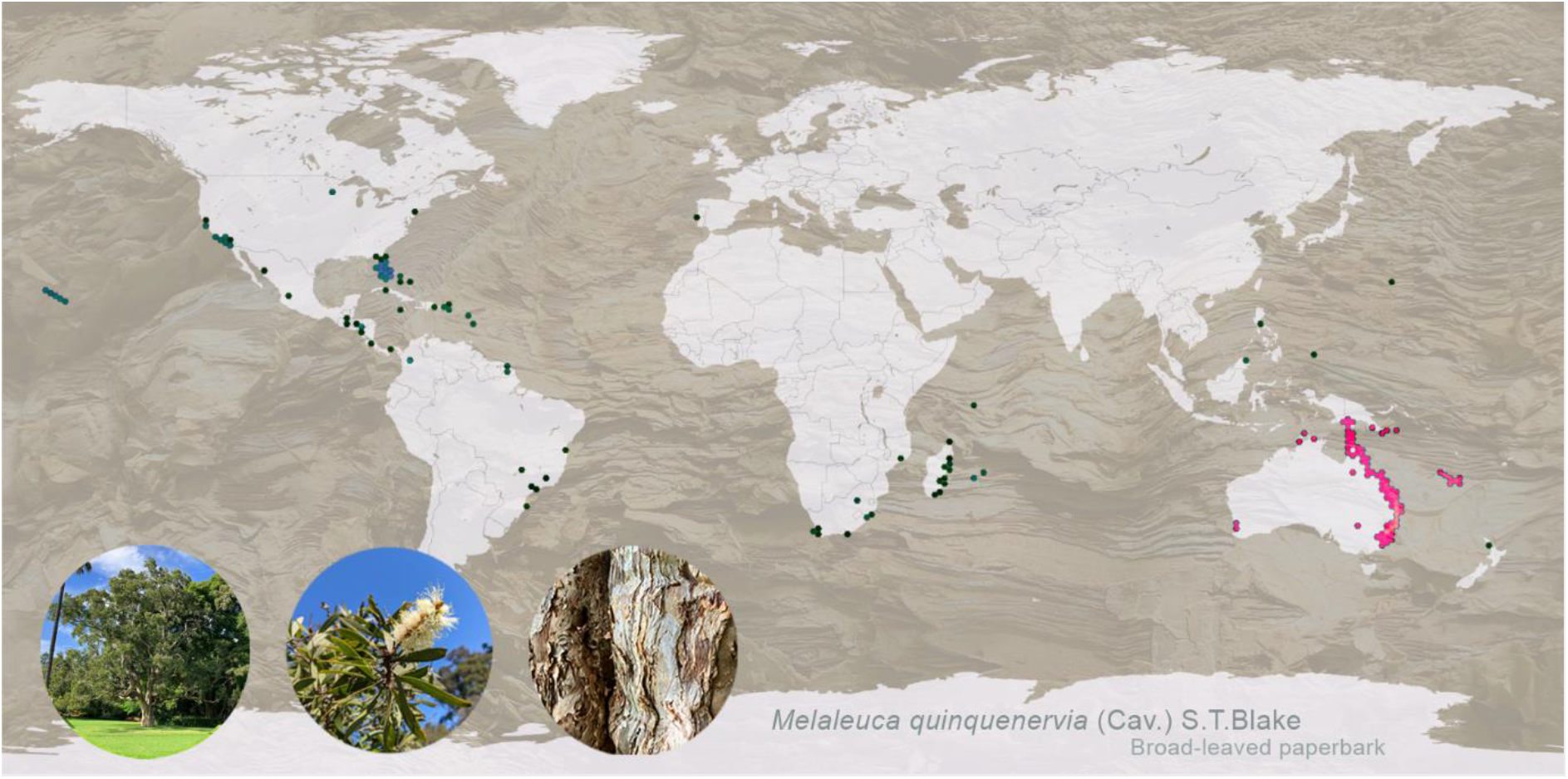
Global distribution of *Melaleuca quinquenervia* in its native range (Australia, Papua New Guinea, New Caledonia and Indonesia; pink dots) and introduced range (blue dots). Data sourced from GBIF with darker shades indicative of higher record densities. Map generated using OpenStreetMap, licensed under the Open Data Commons Open Database License. Photos of the genome tree and detail of bark used in map background taken in the Royal Botanic Garden Sydney by SH Chen and PA Tobias.

Long living tree species, such as *M. quinquenervia*, are exposed to extensive biotic stresses over their lifetime, including a wide range of pests and pathogens. Plants employ various strategies to combat pests and pathogens. These include preformed physical barriers such as leaf cuticles [8,9] and changes in leaf anatomy [10], and chemical barriers such as secondary metabolites [11,12]. At a molecular level, plants rely on an innate immune system to recognise and respond to pathogens [13]. The plant immune system can be considered as two distinctly activated, but interplaying pathways involving cross talk between pathogen and host [14]. Research has therefore focussed on understanding the molecular basis of host tree responses to inform management, with a key emphasis on recognition and response to invasion patterns [15].

There has been substantial research focused on understanding the rapid, cascading response leading to programmed cell death, initiated by resistance receptors of the <u>N</u>ucleotide-binding <u>L</u>eucine-rich <u>R</u>epeat (NLR) domain-type [16]. The genes encoding NLRs are the largest group of plant resistance genes and are modular in their structure, generally containing three main domains: a nucleotide binding (NB) domain, an N-terminal domain, and a C-terminal domain. The NB site, or NBARC (Apaf- 1, R-protein and CED-4) is highly conserved in plants, having an important role in activation of the hypersensitive response (HR) which blocks disease progression by stimulating programmed cell death within and around the infected region [17]. Of the 8 motifs constituting the NBARC, the P-loop motif is the most highly conserved, being essential for ATP hydrolysis and NLR function [18]. The NLR N-terminal domain is commonly a Toll/Interleukin-1 receptor/ Resistance protein (TIR) domain, a coiled-coil (CC) domain, or a RESISTANCE TO POWDERY MILDEW 8-like coiled-coil (RPW8/CC-R) domain [19]. Studies have demonstrated an important role for this domain for pathogen recognition and signalling [20,21]. Plant NLRs also contain leucine rich repeats (LRRs) which are subject to strong diversifying selection and show high sequence diversity even within closely related genes [22]. Studies suggest the high diversity of this region is the result of co-evolution between host and pathogen with several studies showing specific pathogen ligand interaction at this site [22].

While NLRs share common domains, they are highly diverse, even within the well-studied model species *Arabidopsis thaliana* [23]. Adding to this diversity, is the addition of novel integrated domains (IDs) which can be numerous within a NLR protein and are located at various locations within the modular structure of these proteins [24]. Mimicking host proteins, evidence suggests that these domains function as decoy targets for pathogen secreted molecules, known as effectors, allowing for host recognition and triggering immune signalling [25]. A well-documented example is the RRS1 NLR in *A. thaliana* which carries a WRKY domain [26]. It interacts with RPS4 to recognise effectors from a range of pathogens, with the pair forming a complex that is activated upon targeting/modification of the WRKY domain [26]. Without this recognition, pathogen effectors were found to inhibit host WRKY DNA-binding that plays a role in defence signalling, indicating a role for the ID as a decoy [26]. Other notable examples include RGA5 and Pik-1 in rice which both contain a heavy metal associated domain that recognise effectors from the rice blast pathogen *M. oryzae* [27,28].

NLR genes are also known to be numerous in many plant genomes [29], representing over 2% of all genes in apple (*Malus domestica*) [30]. While initial studies computationally identified 149 putative NLR-type genes in the genome of *A. thaliana* [31], more recently, a core set of 106 NLR orthogroups (6,080 genes) has been established across 52 plant accessions largely found in Europe [23] showing the incredible diversity of these genes within a single species. Despite the importance of this gene family in determining plant disease resistance, only 481 genes from 31 species have been fully or partially functionally characterised [32].

Overcoming the challenges associated with assembling these highly polymorphic and repetitive genes has been aided by the latest generation sequencing technologies such as Oxford Nanopore Technologies (ONT) and PacBio HiFi [33,34]. By facilitating the generation of more contiguous genome assemblies, these technologies allow for greater characterisation of, and evolutionary analysis of NLR genes. This was highlighted in recent analysis of an updated reference genome of barley [35] which revealed over double the number of NLR genes compared to previous assemblies generated with short-reads [36,37]. It has also aided in the generation of a near complete NLRome in *A. thaliana,* allowing for the mapping of NLR genes which were previously uncharacterised [23].

The genomes of many diploid organisms are represented as collapsed consensus sequences from homologous chromosomes [38]. Owing to the highly repetitive nature of plant NLRs, detailed genome wide analysis of NLR allelic variation is yet to be carried out. Studies have revealed extensive allelic variation in *NLR* genes such as 8 brown planthopper resistance gene in *Oryza sativa* [38]. These results indicate the importance of detailed analysis of both chromosome sets to more accurately characterise NLRs, with the outcomes having implications for plant:pathogen coevolution and informing downstream molecular analyses. Recent developments in sequencing and scaffolding methods [39] provides the opportunity to generate phased genomes of highly heterozygous organisms such as *M. quinquenervia* [5,40].

Here we present a chromosome-level and pseudo-phased diploid genome assembly for *M. quinquenervia.* We make available FindPlantNLRs [41], a novel pipeline to fully annotate putative NLR genes, taking a genome file as the starting point (Figure 2). We compare NLR allelic variance within the phased, chromosome-level genome assembly of *M. quinquenervia* to provide the first example, to our knowledge, of NLR diversity in a diploid tree genome. Our data indicates that copy number, presence/absence and integrated domains are highly variable between haplotypes. These findings reveal the high level of diversity that exists for *NLRs* within a single plant genome. With much of this lost in a collapsed form, we demonstrate the importance of our approach to assist research into plant responses to environmental challenges.

**Figure 2.**
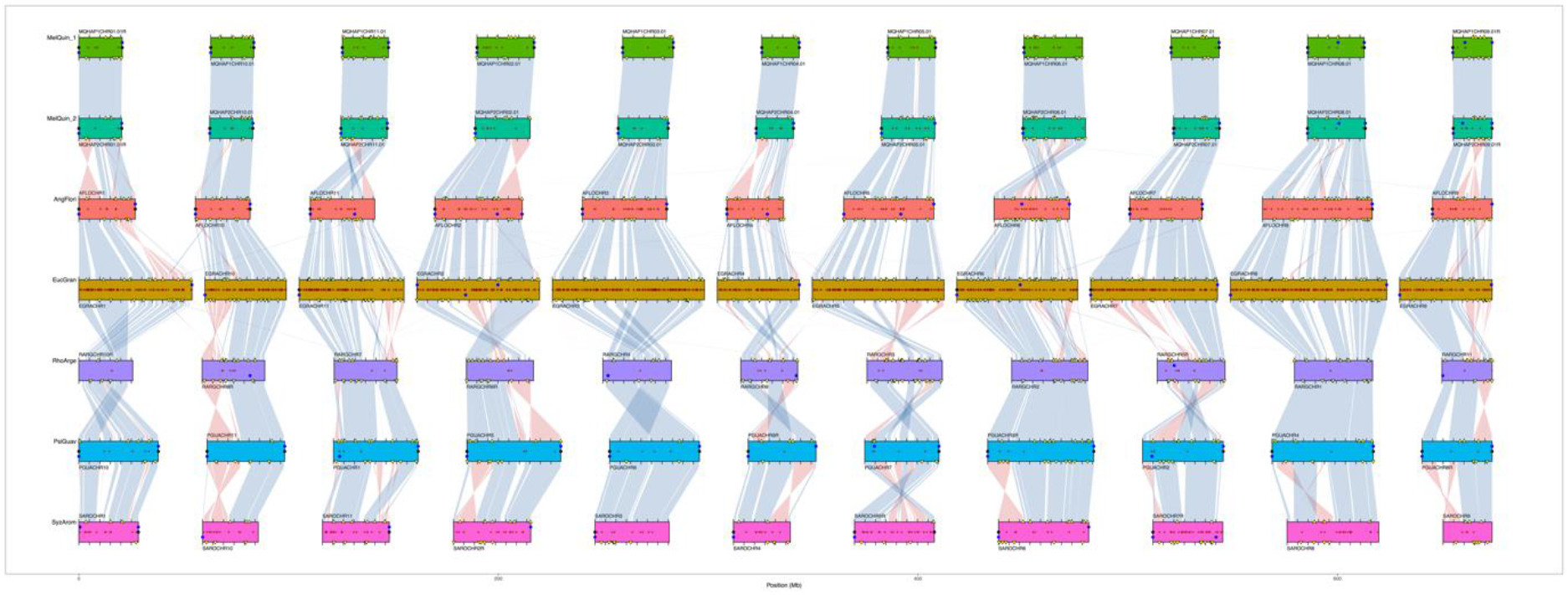
Synteny between *Melaleuca quinquenervia* phased genome and selected chromosome-level Myrtaceae genomes (*Angophora floribunda*, *Eucalyptus grandis, Rhodamnia argentea*, *Psidium guajava* and *Syzygium aromaticum*). Synteny blocks of collinear “Complete” BUSCO genes link scaffolds from adjacent assemblies: blue, same strand; red, inverse strand. Yellow triangles mark “Duplicated” BUSCOs. Filled circles mark telomere predictions from Diploidocus (black) and tidk (blue). Assembly gaps are marked as dark red + signs.

## Analyses

### A high quality pseudo-phased genome assembly for Melaleuca quinquenervia

We sourced leaf material from a mature *M. quinquenervia* tree growing at the Royal Botanic Gardens (RBG) Sydney, New South Wales, for use as the reference genome. The tree was planted in 1880, is 140 years old, of unknown provenance, and is a vouchered specimen of the RBG living collections. High molecular weight DNA was extracted for PacBio HiFi and Oxford Nanopore Technologies (ONT) sequencing. Fresh leaf samples were sent for Hi-C library preparation and sequencing. We assembled the *M. quinquenervia* genome with HiFiasm [42] software using HiFi sequencing data and integrating Hi-C data, with a total yield of 19.46 Gb and 116.4 Gb reads respectively (Table 1). We independently scaffolded the resulting pseudo-phased outputs using the Aidan Lab pipelines [43–45] and determined each haplotype comprised of 11 chromosomes with 94% of sequences assigned to chromosomes for both haplotypes (Figure S1A and B). To independently verify the HiFi assemblies, we assembled and scaffolded the ONT data (Figure S1C and D) which showed a high degree of synteny to the HiFi assemblies (Figure S2A and B). Our final assembly genomes were 269,244,392 bp and 271,680,404 bp for Haplotype A and B respectively (Table 2). We used Chromsyn [46] to investigate synteny of *M. quinquenervia* to five chromosome-level Myrtaceae genomes, all with *2n* = 22 chromosomes (Figure 2). The scaffolding of Haplotype A is supported by the scaffolding of Haplotype B, for *M. quinquenervia*, despite the processes being run independently. We determined some inversions against the other Myrtaceae genome chromosomes that likely represent misassemblies in the less contiguous assemblies (Figure 2).

**Table 1.** Genomic sequence reads for the *Melaleuca quinquenervia* genome.

| Sequencing platform | Library | Median insert size (bp) | Mean read length (bp) | No. of reads | Sequence bases (Gb) |
| --- | --- | --- | --- | --- | --- |
| PacBio Sequel II | HiFi SMRTbell | 16,506 | 17,058 | 1,140,849 | 19.46 |
| Illumina NextSeq 500<sup>‡</sup> | Phase Genomics<br>Proximo Hi-C (Plant) | - | 2 x 151 | 770,901,164 | 116.4 |
| Oxford Nanopore Technologies | Ligation (SQK-LSK110) | - | 26,803 | 2,400,431 | 64.68 |
| **Total gDNA** | - | - | - | **774,442,444** | **200.5** |

<sup>‡</sup> Includes a pilot iSeq run used to QC the library

**Table 2.**
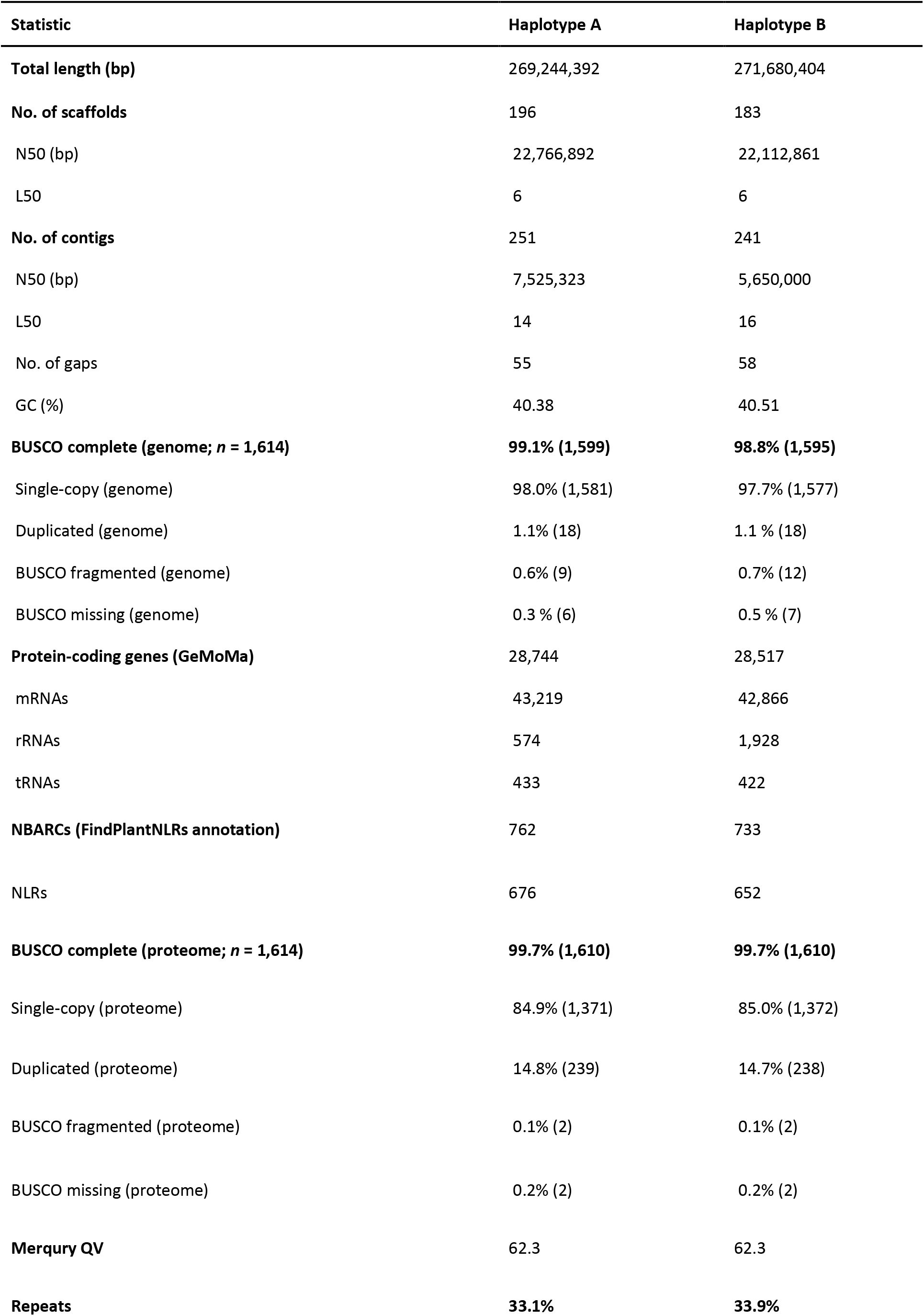
Genome statistics for the *Melaleuca quinquenervia* phased reference genome.

We checked the genome outputs using Depthsizer [47] using HiFi and ONT reads to show a genome size of approx. 274 Mbp and 272 Mbp for Haplotype A and B, respectively, with the ONT assembly giving similar figures (Table S1). We further validated the genome size using GenomeScope [48] which predicted a haploid genome size of 262 Mbp (Figure S3A). We confirmed the diploid state of the genome using SmudgePlot [49] (Figure S3B).

To improve the overall quality of the *M. quinquenervia* genomes, we carried out several rounds of scaffolding, polishing and gap filling, with telomeres predicted by both Diploidocus [47] and tidk [50] at the end of chromosome scaffolds in most instances (Figure S2A and B). There are only a small number of gaps (fewer than 60) (Figure S2A and B).

Base pair level accuracy was tested against Merqury [51]with both haplotypes showing very high quality and accuracy scores. Additionally, we determined very high genomes completeness of both haplotypes using Benchmarking Universal Single Copy Orthologs (BUSCO) [52] (Table 2, Figure 3A and B, Figure S4A-F). We ran GeMoMa [53] annotation on the two haplotypes and both proteomes were 99.7% complete according to BUSCO. We assessed the repetitive, as well and transfer (tRNA) and ribosomal RNA (rRNA) elements using RepeatModeler [54] (Table 2).

**Figure 3.**
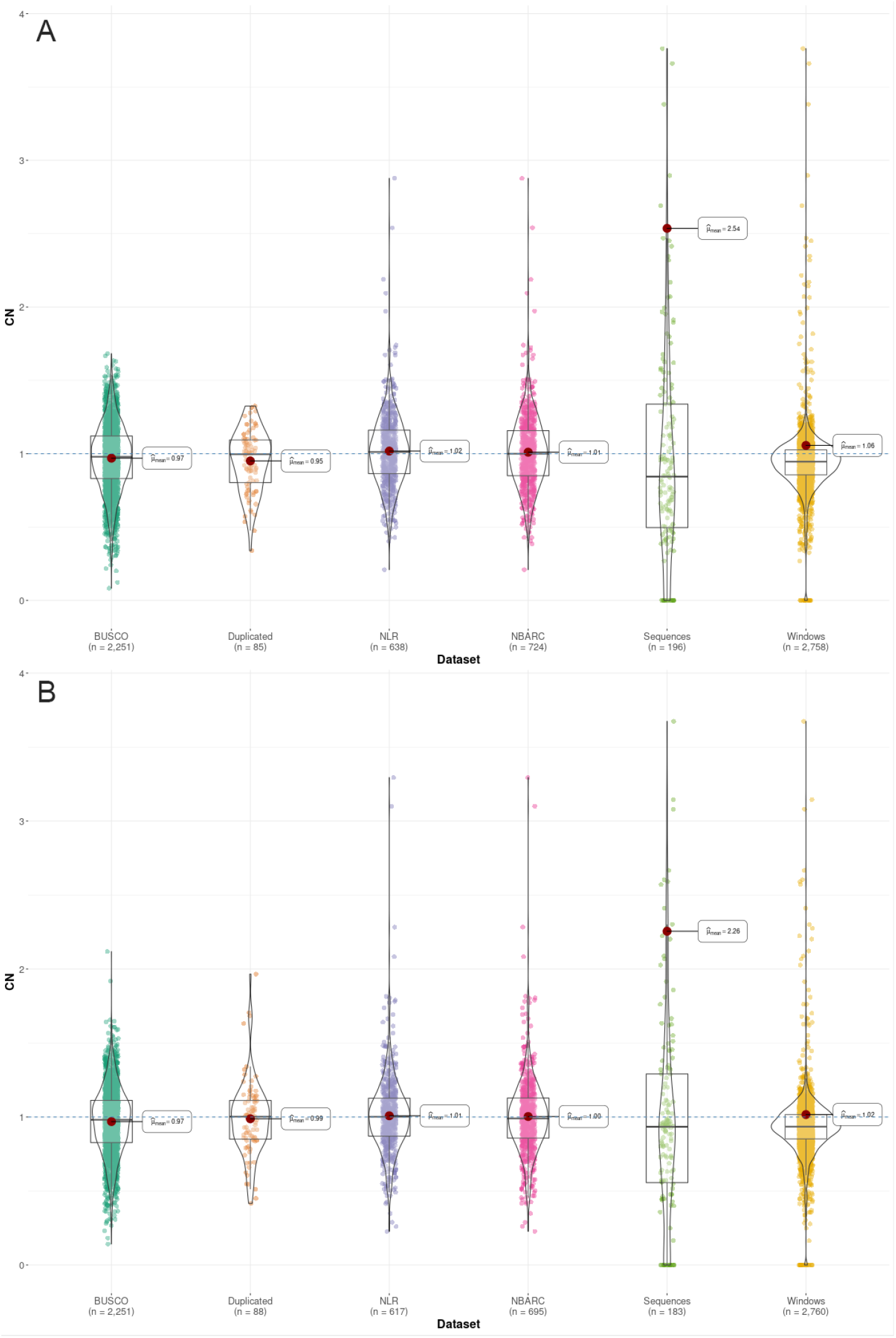
Genome-wide regional copy number analysis for *Melaleuca quinquenervia* (A) Haplotype A and (B) Haplotype B using HiFi read data. Copy number (CN) is relative to a single diploid (2*n*) copy in the genome. Violin plots and means generated with ggstatsplot. Each data point represents a different genomic region: BUSCO, BUSCO v5 (MetaEuk) single-copy “Complete” genes; Duplicated, BUSCO v5 “Duplicated” genes; NLR, resistance gene annotations; NBARC, NBARC domains; Sequences, assembly scaffolds; and Windows, 100 kb non- overlapping windows across the genome. Plot truncated at CN = 4.

**Figure 4.**
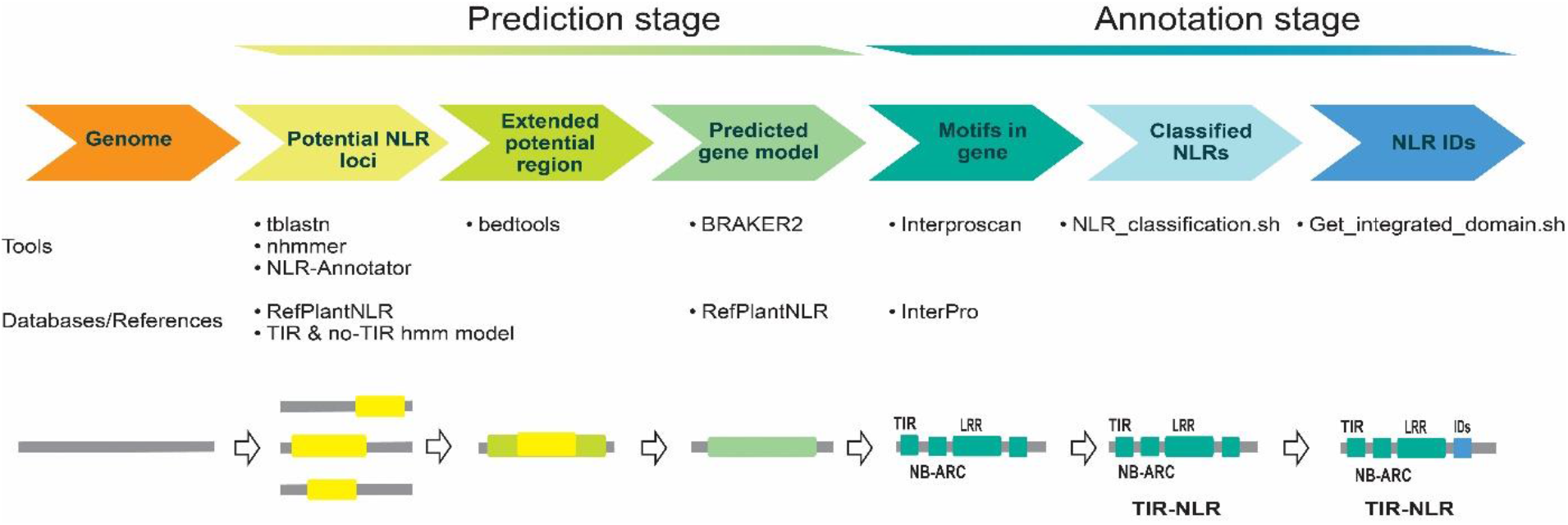
Workflow of the FindPlantNLRs pipeline: a tool for annotating nucleotide-binding and leucine-rich repeat (NLR) genes. The pipeline annotates predicted NLR genes from an unmasked genome fasta file input. We combine loci identified using NLR-annotator software with a basic local alignment search tool (tblastn) using recently compiled and functionally validated NLR amino acid sequences and a nucleotide iterative Hidden Markov Model (HMM) [95] to locate NBARC domains in genomes. The loci identified (including 20 kb flanking regions) are then annotated with Braker2 software using protein hints from experimentally validated resistance genes. Annotated amino acid fasta files are screened for domains using Interproscan and the predicted coding and amino acid sequences containing both NB-ARC and LRR domains are located back to scaffolds and extracted in gff3 format.

### A novel pipeline to identify and classify NLRs

We developed a comprehensive pipeline to annotate predicted NLR genes from an unmasked genome fasta file input. The rationale for an unmasked sequence is that the repetitive nature of the NLRs, regions may be missed with standard annotations. Our pipeline, named FindPlantNLRs [41] utilises three key approaches. We combined loci identified using (1) NLR-annotator software [55] with (2) a basic local alignment search tool (tblastn) [56] using recently compiled and functionally validated NLR amino acid sequences and (3) a nucleotide iterative Hidden Markov Model (HMM) [57] to locate NBARC domains in genomes [58,59]. While the pipeline was developed to seek NLR genes within Myrtaceae genomes, the supplied NBARC HMMs are suitable for any plant genome search due to the iterative step that builds a unique species-specific HMM combined with the use of two other steps that incorporate broader models. The loci identified through these methods, and including 20 kb flanking regions, are then annotated with Braker2 software [60] using protein hints from experimentally validated resistance genes [32]. Annotated amino acid fasta files are screened for domains using Interproscan [61] and the predicted coding and amino acid sequences containing both NBARC and LRR domains are located back to scaffolds and extracted in additional scripts are available on GitHub. To identify all classes of annotated NLRs, we developed a script that sorted and classified the “gene” types. We ran the file outputs from FindPlantNLRs with the NLR classification script [41]. To further identify novel predicted integrated domains in the annotated NLRs, we developed a script to search the data based on PFAM domain identities not classically associated with NLRs [41].

### NLR number is variable across chromosomes and haplotypes

Using the FindPlantNLRs pipeline, we identified 762 putative NBARC containing genes in Haplotype A and 733 in Haplotype B based on the presence of the NBARC domain (Table S2). As NLRs require both NBARC and LRR regions to be functional, for downstream analyses we were interested in isolating full gene models (genes containing both domains). Termed NLRs from hereon, we have divided these into genes containing a TIR domain (TNL), a CC or Rx domain (CNL), and those lacking TIR or CC domains (NL). Of the 762 NBARC containing genes in Haplotype A, we predicted 676 NLRs of which 67 lacked an N-terminal CC or TIR domain (Table S3). We excluded 86 predicted genes as they did not fit the definition of full genes models, with 68 lacking a C-terminal LRR domain and 18 lacking both N and C terminal domains (Table S2). Of the 733 NBARC containing genes in Haplotype B, we predicted 652 full gene models of which 71 lacked an N-terminal CC or TIR domain (Table S3). We excluded 81 predicted genes as they did not fit the definition of full genes models, with 61 lacking a C-terminal LRR domain and 20 lacking both N and C terminal domains (Table S2).

As NLR numbers differed between haplotypes, we sought to further investigate this difference at the chromosome level. The number of genes per chromosome varied by up to 31 genes between haplotypes, with only chromosomes 1 and 9 containing the same number of genes across Haplotypes (Figure 5A). In Haplotype A, chromosomes 2 contained the highest number of NLR genes followed by chromosomes 5 and 3 while chromosome 5 contained the highest number of genes followed by chromosomes 3 and 2 in Haplotype B (Figure 5A). Upon further investigation, we determined the classes of NLRs is also consistent across chromosomes 1 and 9, while all other chromosomes the number of NLRs in each class is variable. (Figure 5B and C). Chromosome 1 was also the only chromosome to contain NLRs of one class (CNL) (Figure 5B and C).

**Figure 5.**
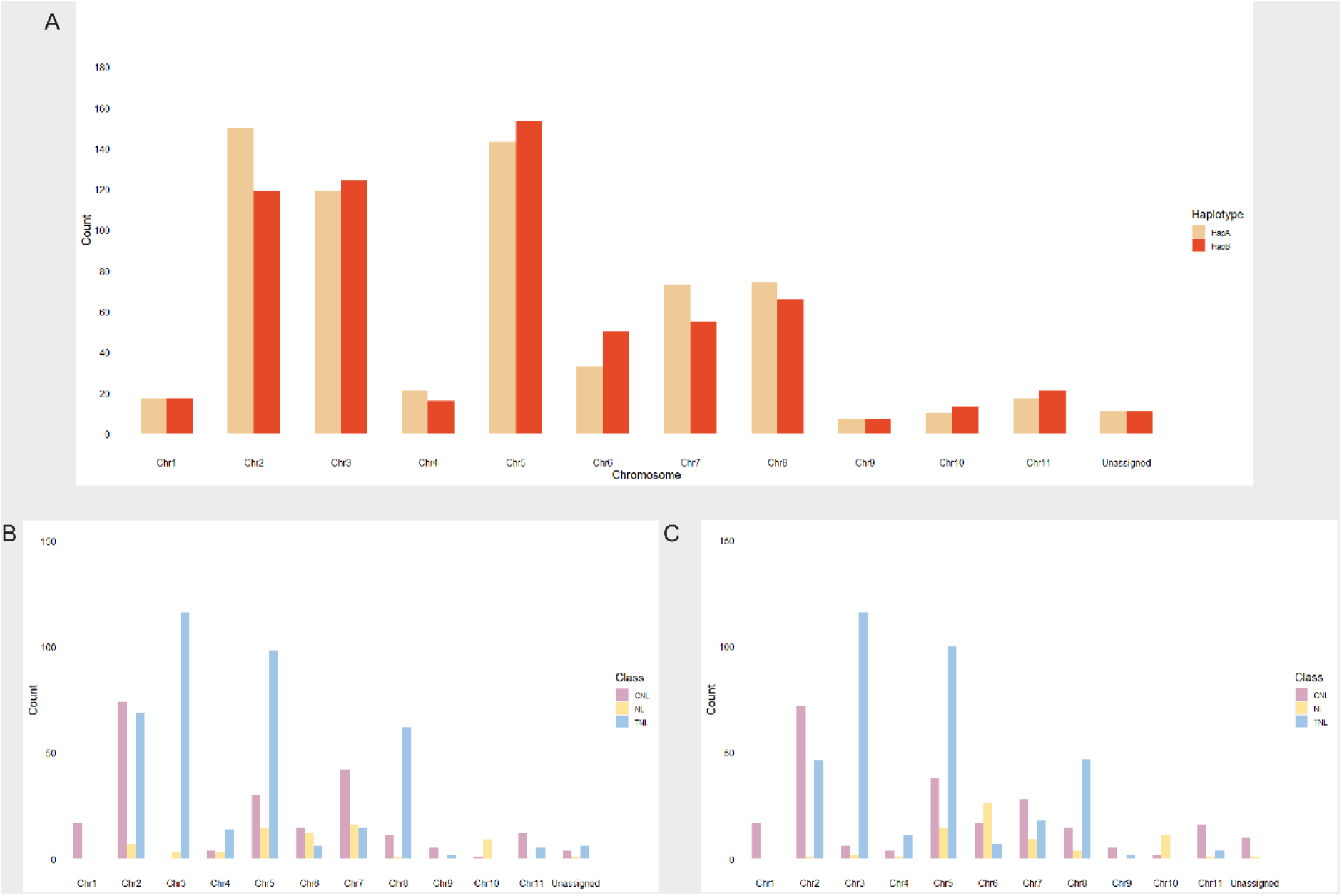
Summary of the number of predicted NLR genes per chromosome in the phased *Melaleuca quinquenervia* genome. (A) Comparison of the number of putative NLR genes on each chromosome in Haplotypes A and B. Putative NLRs were classified into TIR-NLR (TNL), CC-NLR and Rx-NLR (CNL) and NL classes on individual chromosomes in (B) Haplotype A and (C) Haplotype B.

### NLR genes are arranged in clusters with hotspots on chromosomes

To visualise the physical clustering of NLRs on chromosomes, we mapped gene locations to chromosomal locations in both Haplotypes (Figure 6A and B). Employing the definition of a cluster as being a genomic region with 3 NLRs less than 250 kb apart with fewer than 8 other genes between each NLR, we determined variation in the number of genes clustering per haplotype, and clusters per chromosome within and between haplotypes. At a gene level, we determined 89.8% of genes in Haplotype A and 90.5% of genes in Haplotype B occur in clusters. A total of 51 clusters were identified in Haplotype A with an average of 4.6 clusters per chromosome and an average of 11.7 genes per cluster. A total of 50 clusters were identified in Haplotype B, averaging 5 clusters per chromosome and an average of 11.4 genes per cluster. 5.1% of genes were determined to occur as singles in Haplotype A and 5.1% as pairs. 6.1% of genes in Haplotype B were determined occur as singles and 3.4% as pairs. In both haplotypes, the most clusters were on chromosome 5 (11 and 15 on Haplotypes A and B respectively) and the least (one cluster) on chromosome 9 in both Haplotypes (Figure 8A and B). The independently assembled and annotated assemblies based on ONT data verified the location of the majority of NLRs (Figure S5).

**Figure 6.**
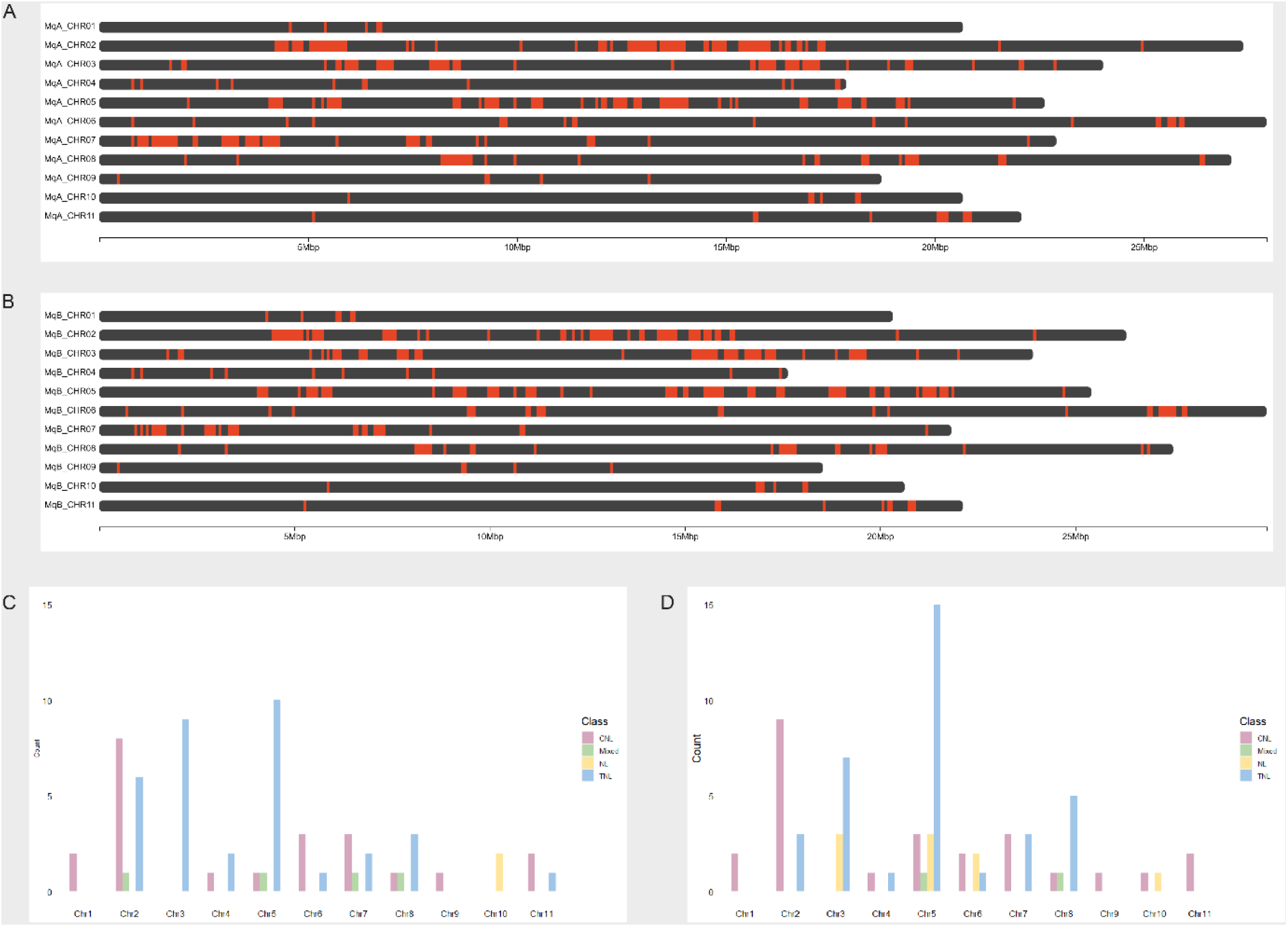
Physical clustering of predicted NLR genes in the phased *Melaleuca quinquenervia* genome. Physical locations of predicted NLR genes on the chromosomes of *Melaleuca quinquenervia* (A) Haplotype A and (B) Haplotype B generated using ChromoMap in RStudio. The number of clusters per chromosomes in (D) Haplotype A and (E) Haplotype B was analysed and categorised based on the classes of all NLR genes.

We then determined if these clusters were comprised of genes of the same class. We defined classes of clusters by clusters containing only genes of one class along with *NL*-type genes, otherwise they are considered mixed. TNL-type clusters were the most abundant clusters in both haplotypes and most abundant on chromosomes 3 and 5 in Haplotype A and chromosome 5 in Haplotype B (Figure 6C and D). CNL-type clusters were more evenly distributed across chromosomes in both haplotypes, with chromosome 2 containing the most clusters (4 in Haplotype A and 5 in Haplotype B) (Figure 6C and D).

### Integrated domains are unique between haplotypes

Based on PFAM domain identities of the predicted NLR genes, we discovered 4.8% of NLRs in Haplotype A contain novel integrated domains (IDs) (Figure 7A), of which 46.9% contain more than one domain. Similarly, we observed a comparable percentage of 4.5% in Haplotype B (Figure 7B), with 51.7% of the predicted genes containing multiple domains. We also examined the number of ID-containing NLRs per chromosome and noted that in Haplotype A, chromosome 3 had the highest count with seven while chromosome 11 had none. In Haplotype B, chromosome 3 had six ID-containing NLRs, and 11 also had none (Figure 7C). During our investigation, we identified 48 unique IDs across both haplotypes. Interestingly, we found 23 IDs were exclusive to Haplotype A but only eight were exclusive to Haplotype B (Table S4). The remaining IDs were identified in both haplotypes (Table S4).

**Figure 7.**
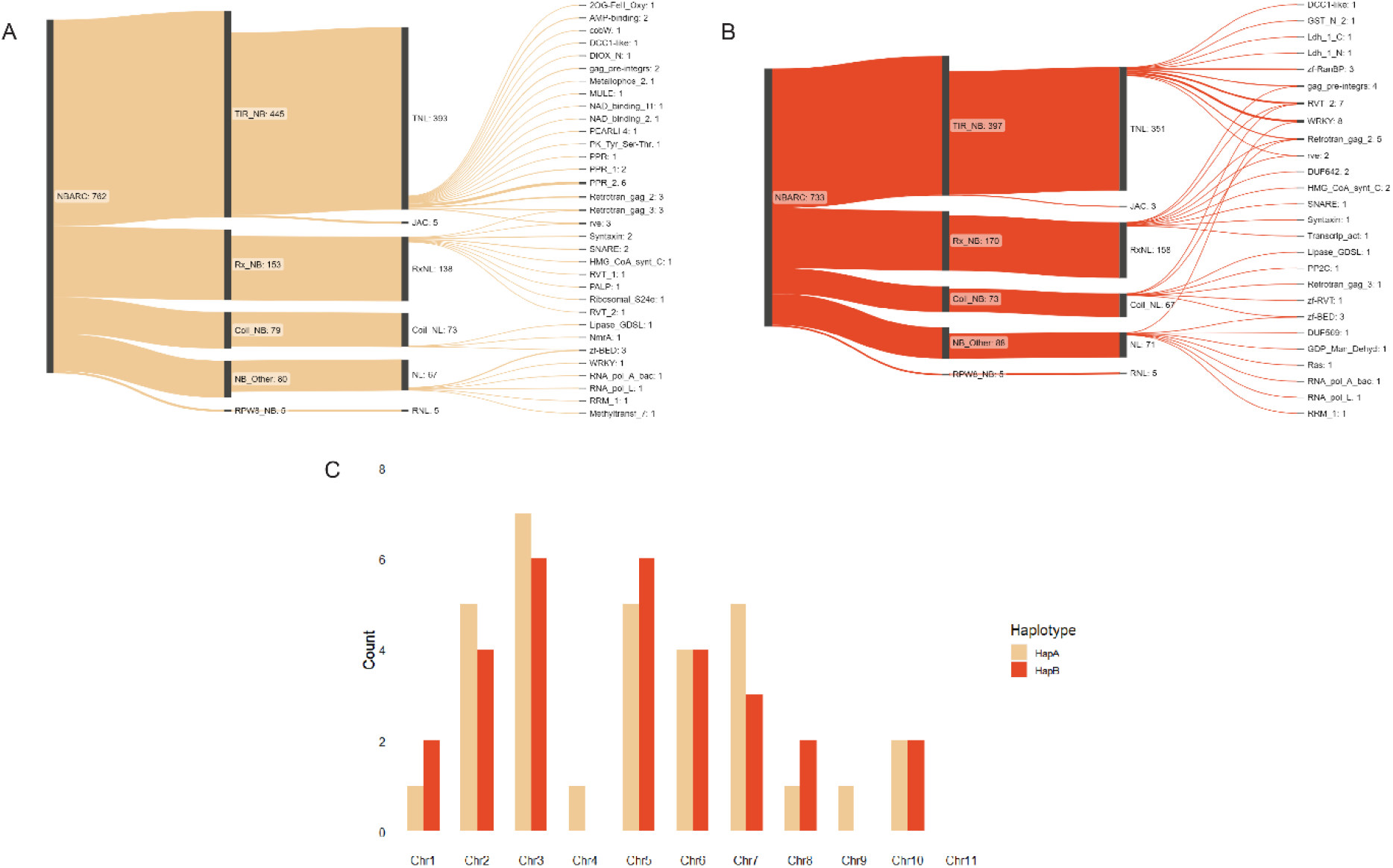
The NLR gene complement in the phased *Melaleuca quinquenervia* genome. The two sets of chromosomes corresponding to (A) Haplotypes A and (B) B were independently classified and visualised to present the domain classes using Sankeymatic [62] including novel integrated domains (IDs) with abbreviations derived from Pfam database (REF). NB = Nucleotide Binding Domain, TIR = Toll/Interleukin-1 receptor, JAC = Jacalin Domain, Rx = Potato CC-NB-LRR protein Rx, Coil=Coil-Coil Domain, RPW8 = RESISTANCE TO POWDERY MILDEW 8-like coiled-coil (C) The number of ID-containing NLRs per haplotype and chromosome in both haplotypes.

**Figure 8.**
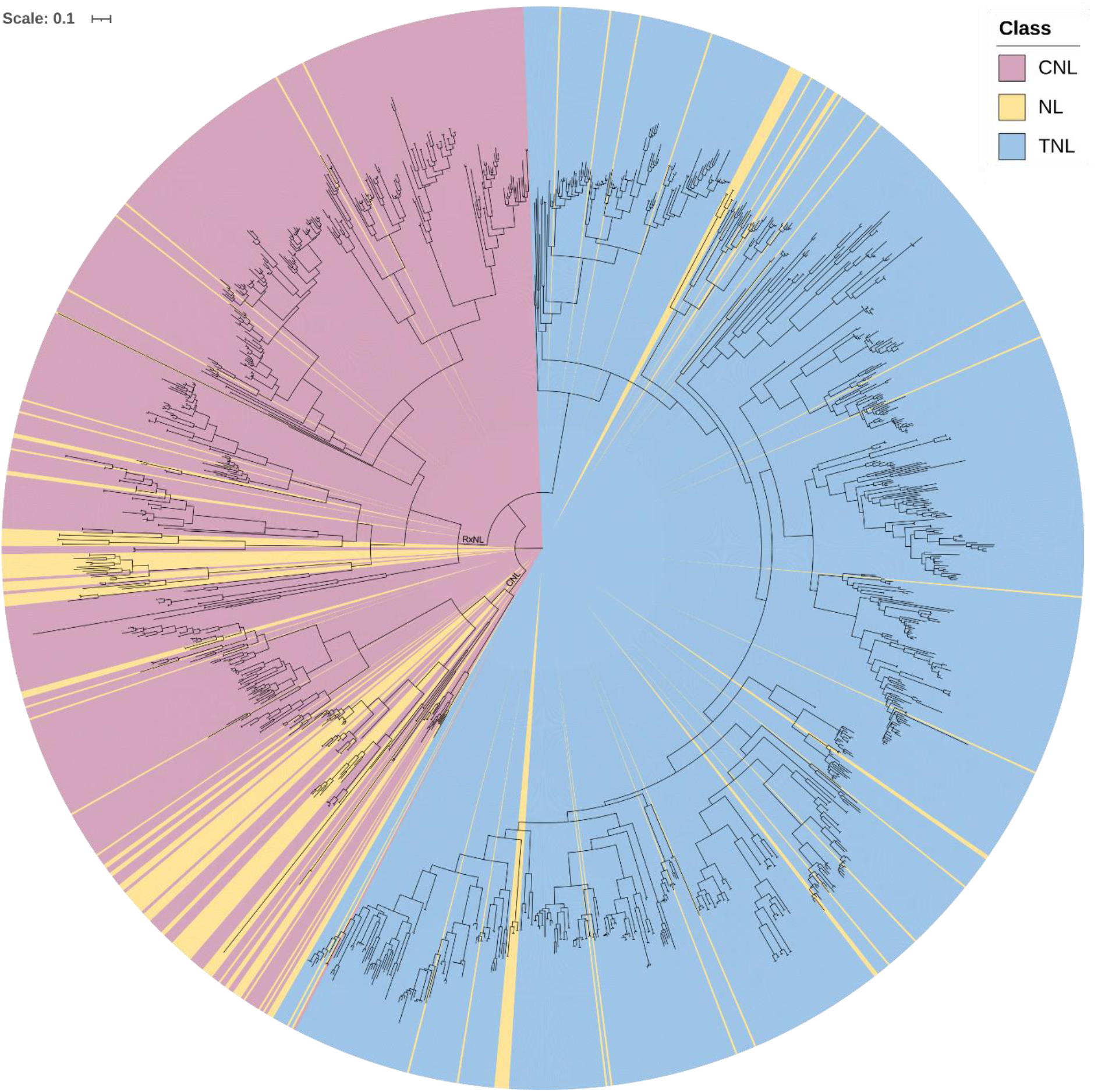
Evolutionary relationship of NBARC domains from predicted NLR genes within the phased *Melaleuca quinquenervia* genome. The NBARC domain fasta file and additional NBARC sequences, as outgroups, from functionally validated plant NLRs [32], were aligned with clustal-omega (v.1.2.4). The phylogenetic tree was inferred with the alignment file using iqtree (v.1.6.7) and visualised in iTOL (v.5). Each tip represents one putative *NLR* gene with branch lengths signifying rates of amino acid substitutions. Colours indicate the CNL (including RxNLs) (pink), TNL (blue) and NL (yellow) clades. Scale = 0.1 amino acid substitutions per site.

### NLRs cluster into two distinct clades

The evolutionary relatedness of the 1,328 NBARC domains (462 CNL, 726 TNL, and 140 NL) from complete NLR genes models separated into two major clades: CNL (CNL, RxNL and RNL genes combined) and TNL genes (Figure 8). Fifty-nine percent of all sequences aligned with the TNL (784) clade and forty-one percent of total sequences aligned with the *CNL* clade (544) with 98 of the 140 NL sequences aligned with CNL and 42 aligned with TNL clades (Figure 8). Fifteen *CNL* NBARC sequences clustered within the TNL clade, however no TNLs clustered within the CNL clade. On closer inspection of these fifteen NBARC amino acid sequences, we determined that the integrity of the tree is correct due to the lack of the ‘W’ (tryptophan) at the ‘LDD*W’ kinase 2 sub-domain (Figure S6). This is canonical for CNL clade NBARC domains but not present in TNL clade [59]. We inspected the annotation and classification from FindPlantNLRs and found coiled-coil and Rx domains at the amino-terminus on these fifteen gene models, hence the classification. It should be noted that all other NLR analyses in our study are based on the full annotated gene classification.

### Transcript evidence found for predicted NLRs

To confirm that in-silico NLR predictions were actively expressed, we downloaded RNAseq data from a previous *M. quinquennia* study that investigated responses to the plant pathogen causing myrtle rust [63]. We mapped all the available RNASeq data to the NLR coding sequencing for each haploid genome independently using Hisat2 [64]. Taking the transcripts per million (TPM) cut-off of 50, we determined expression for 617 and 596 NLR coding sequences from Haplotype A and B respectively. The most abundantly expressed predicted *NLR* gene is an *RPW8* (PF05659) NLR homologue, TPM 50,744 and 47,856 for Haplotype A and B respectively. This gene is predicted on chromosome 6, NLR gene identifications, g7145.t1 and g1651.t1 respectively (Table S3).

## Discussion

### A high-quality diploid genome for the keystone wetland species, Melaleuca quinquenervia

To promote scientific investigation, we have assembled a telomere-to-telomere diploid genome for a keystone wetland species, the broadleaved paperbark tree, *M. quinquenervia*. Using ∼70x HiFi coverage (35x per haplotype), combined with ∼380x Illumina Hi-C coverage, our assembly scaffolded into the expected eleven Myrtaceae chromosomes (*2n* = 22) and has a very high level of BUSCO completeness (Table 2). With careful curation to remove scaffolding errors and misassemblies, followed by polishing, we numbered two sets of parental chromosomes in accordance with the Myrtaceae reference genome, an inbred clone of *Eucalyptus grandis* [6]. We were able to show synteny between the *M. quinquenervia* chromosomes and with five other publicly available chromosome-level Myrtaceae genomes (Figure 2). Additionally, the genome and subsequent analyses were independently validated with scaffolded assemblies using ∼234x ONT data. Based on homology with three publicly available Myrtaceae proteomes and with *A. thaliana*, we predicted 28,744 and 28,517 protein coding genes within the two chromosome sets. These numbers are slightly less per haplotype, but comparable to the predicted 36,779 for the haploid genome of *E. grandis*. This is likely to be due to the earlier generation sequencing technology, assembly software and the result of collapsed assemblies for highly heterozygous plants. We annotated repetitive genomic regions at ∼33 percent in both haplotypes, compared to 41 and 44 percent in *E. grandis* [6]and *E. pauciflora* [65] respectively, likely related to the smaller genome size for *M. quinquenervia*. There was a marked difference in rRNA content between the two haplotypes and these differences are being driven by rRNA on unanchored contigs. Our curated assembly meets the high standards and metrics of the vertebrate genome project objectives [66] providing an exceptional resource for functional molecular and evolutionary studies.

### A smaller than predicted genome for Melaleuca quinquenervia

A 2C-value of 1.94 was previously reported in the literature using flow cytometry on samples from a tree in a university garden [66]. We therefore expected the genome size for each haploid assembly to be 949 Mbp and planned our sequencing experiments accordingly. The *M. quinquenervia* genomes we assembled are much smaller, at ∼270 Mbp, and polyploidy has not been reported in this species. The authors on the flow cytometry study reported problems processing their Myrtaceae samples, perhaps explaining the large size discrepancy in these results. To test that our results were accurate, we checked the ploidy and ran kmer- and read depth-based analyses, as described in the methods. Results indicated the genome was 270-280 Mbp, less than half the size of the *E. grandis* genome at 640 Mbp [6]. While the genome size was surprising, we were able to use the high sequence coverage to ensure a highly accurate diploid genome.

### The annotated NLR complement for both Melaleuca quinquenervia chromosome sets

With the high quality of our genome, we were able to comprehensively annotate the NLR-type resistance genes in both inherited chromosome sets, using our novel FindPlantNLRs pipeline. Of the 1,495 annotated NBARC containing genes identified in the *M*. *quinquenervia* genome (Figure 5), we determined that 1,328 were complete NLRs while a further 167 contained the NBARC domain but lacked either, or both, the C or N-terminal domains. The number of NBARC containing genes in the genome is consistent with analysis of *E. grandis* which was determined at 1487 NBARC containing genes [59] despite a much larger genome size. Although genome size is not directly correlated with NLR content [67], the presentation of *E. grandis* genome in its collapsed form may result in underrepresentation of the NLRs as allelic variants. We estimated up to 52 genes had no ortholog in either haplotype, while up to 73 only contained ortholog in the corresponding haplotype (Table S6). To our knowledge, this is the first published research that has presented the allelic NLR complement in a phased, chromosome-level genome. As such, analysis of ortholog between haplotypes is limited to software which compares individual species which may lead to the discrepancies in ortholog numbers in our analyses (Table S6). Nonetheless, our detailed analysis highlights unique allelic variation that will assist research into the reported different phenotypic responses to pest and pathogen challenged with the family Myrtaceae [63]. Our data might also be useful for understanding the strong evolutionary selection pressures on these plant immune receptors that has resulted in the allelic variation we present for *M. quinquenervia*.

### Melaleuca quinquenervia NLRs are dominated by TNL-type resistance genes

Consistent with the *E. grandis* NLR annotation, is the higher proportion of TNL to CNL type genes supporting an expansion of the TNL clade within the Myrtaceae [59]. This is further validated by recent phylogenetic analyses using transcripts from *M. quinquenervia* and *M. alternifolia* which revealed approximately two thirds of NLR transcripts clustering with TNLs from *E. grandis* [68]. We found TNL to CNL ratios of ∼3:1 in Haplotype A and ∼3:2 in Haplotype B of *M. quinquenervia*. The ID containing NLRs had a greater proportion of TNLs than CNLs with IDs (∼2:1 and 3:1 in Haplotypes A and B respectively). The TIR domain has been demonstrated to play a key role in the self-association of the NLR proteins to form higher order resistosomes which are necessary for immune signalling [69]. The expansion of the TIR class combined with fused IDs within TNLs, discussed later, may provide novel defence capacity against pests and pathogens.

### Phylogenetic evolutionary analysis supports the NLR classification results

By combining all the NBARC amino acid domains from both haplotypes, we visualised the evolutionary relatedness of NLRs. While the phylogenetic tree was based of alignment of NBARC domains, and not full annotated genes, it demonstrated the clear divergence into CNL and TNL clades (Figure 8) as observed in other plant species [31,59]. Of the NLRs lacking CC or TIR domains (NLs), 42 are clustered in the TNL clade and the remaining 96 into the CNL clade. Of interest, the expansion of the TNL clade, also observed in *E. grandis* [59] with 53 percent TNL to 47 percent CNL, was comparable in *M. quinquenervia* with 59 percent TNL to 41 percent CNL (Figure 8). There were 15 predicted CNLs that clustered within the TNL clade. On inspection of these amino acid sequences, we found that they had coiled-coil or Rx-type domains fused to classic TNL-type NBARC domains. These might indicate amino terminal domain swapping and might be an evolutionary mechanism, however further functional and molecular validation is required.

### Physical clusters of NLRs on M. quinquenervia chromosomes

Analysis of the putative TNLs, CNLs and NLs within the phased genome of *M. quinquenervia* revealed the majority of NLRs located within clusters, with 86% clustering in Haplotype A and 88% in Haplotype B. Only 14% and 12% from Haplotype A and B respectively did not fall into clusters, compared to approximately a quarter of *NLRs* in *E. grandis* [59], cultivated rice (*Oryza sativa*) [70], and *A. thaliana* [31]. For *M. quinquenervia*, there were approximately 5 NLR genes for every Mbp of the total genome size while in *A. thaliana*, *E. grandis* and *O. sativa* the number of NLRs per Mbp ranged from 1.2 to 2.3 [23,59,71]. The higher density of NLRs in the *M. quinquenervia* genome may explain the higher proportion of NLRs appearing in clusters.

Most clusters were homogenous, containing NLRs of the same class, with only 4 heterogenous clusters in Haplotype A and 2 in Haplotype B (Figure 8D and E). The high proportion of homogenous clusters suggests the expansion of these genes into clusters is driven by tandem duplication [72], and serves as a mechanism for maintaining NLR diversity [73]. Clustering may also play an important role in pathogen resistance. NLR pairs such as *RGA4* and *RGA5* [74] and *Pik-1* and *Pik-2* in cultivated rice [75] are oriented in a head-to-head manner, function cooperatively in pathogen recognition and response, with one acting as sensor of the pathogen and the other as an executor of immune signalling. This was also observed for the NLR pair *RPS4* and *RRS1* in *A. thaliana*, suggesting a shared promoter for the co-regulation of the two genes [76,77]. Interestingly, for each of these pairs, one partner from each contained an ID. On chromosome 3 of Haplotype B of *M*. *quinquenervia*, one pair of NLRs was identified in this head-to-head manner, with one partner containing one RVT2 and one gag_pre- integrs ID. The identification of genes in the head-to-head manner in *M. quinquenervia* may indicate a functional role for these genes in disease resistance, with further studies needed to elucidate a potential function.

### The NLR repertoire is unique between haplotypes

Overall, the patterns of individual NLR numbers, classes, clusters, and cluster types across chromosomes appear consistent between the two haplotypes of *M. quinquenervia* (Figure 5 and Figure 6). However, analysis at the individual chromosome and gene level revealed diversity in the number and classes of genes between haplotypes for all except chromosomes 1 and 9 (Figure 5). While consistent in gene number, and gene number per class, analysis of the IDs across chromosome 1 revealed one gene on Haplotype B to contain two DUF642 domains which was not present on the corresponding gene in Haplotype A. Similarly, one gene in Haplotype A of chromosome 9 contained one NAD_binding_11 and one NAD_binding_2 domains which were not present in the corresponding gene on Haplotype B (Table S3). The presence/absence NLR polymorphisms between the haplotypes of *M. quinquenervia* are likely explained by the outcrossing nature of the species. High levels of genetic diversity maintained in long-lived, outcrossing woody species [78], combined with exposure to a range of pests and pathogens over their lifetime, may lead to changes in NLRs arrangement over subsequent generations. Presence/absence polymorphisms of NLRs has been observed in several plant species such as between inbred accessions of *O. sativa* and *A. thaliana* [79,80]. This may be explained by the fitness cost associated with the maintenance of these genes [81], leading to loss of corresponding genes in the absence of the pathogen.

We identified a total of 53 unique IDs across both haplotypes, accounting for 4.4% of NLR genes in Haplotype A and 6.8 % in Haplotype B. These figures are consistent with a recent review of published NLR-ID analyses that revealed 3.5 – 14% of NLRs contained IDs [25]. These fused integrated domains appear to mimic host proteins that are targets for pathogen effectors, lead to the triggering of defence response [24]. Some of the most commonly occurring integrated domains belong to families of proteins with critical roles in plant defence [24,82] such as WRKY transcription factors and BED zinc fingers (BEAF and DREF from *Drosophila melanogaster* peptide; zf-BED). In the genome of *M. quinquenervia*, one of the most commonly occurring ID was the WRKY domain which was identified in five genes across the two haplotypes. A notable example of the role of an integrated WRKY domain present in an NLR, is the *Arabidopsis Ralstonia solanacearum gene 1* (*RSS1-R*; Le Roux et al., 2015; Sarris et al., 2015). Bacterial effectors were found to bind to the WRKY domain of the NLR protein and other WRKY containing proteins [83], suggesting a role for this domain as a decoy. Another common domain was the zf-BED domain which was identified in seven genes across the two haplotypes. While the function of the ID is yet to be elucidated, zf-BED domains have been observed in NLR genes conferring resistance to rust pathogens in barley and wheat [84,85]. The identification of these fused domains suggests a role for these genes in pathogen recognition.

### Potential implications

Long-lived tree species must respond to a wide range of biotic stresses. Our results provide insight into the diversity of the NLR gene family within a single host tree species, indicating a potential mechanism for responses to invasive pathogens over a life-span. We provide a framework for studying highly repetitive resistance genes by generating a high-quality pseudo-phased reference genome. With advances in sequencing and software, we are beginning to investigate the full repertoire of all genes, including NLRs, here starting with a representative Myrtaceae tree, *Melaleuca quinquenervia*. Given the diversity of NLRs from just two haplotypes, our results indicate that association studies of will need to model presence/absence of NLRs, in addition to segregating sequence variants. Future studies may expand to comparing population level diversity of NLRs and the diversity of NLRomes across woody plants.

## Methods

### DNA extraction and sequencing

#### Sampling and DNA extraction

We obtained young fresh leaves (approximately 30 g) from a mature *Melaleuca quinquenervia* (Cav.) S.T. Blake tree growing at the Royal Botanic Gardens (RBG) Sydney, New South Wales (BioSample accession SAMN20854364) for use as the reference genome individual. We chose this specimen for the ease of ongoing access to leaf, cuttings, and seed material. The tree was planted in 1880 by HRH Prince George of Wales, later King George V. The tree is now 140 years old, of unknown provenance, and is showing signs of senescence.

For PacBio HiFi sequencing, we extracted high molecular weight (HMW) genomic DNA (gDNA) using two sorbitol washes [86] followed by a CTAB/NaCl/Proteinase K protocol [87]. We purified gDNA with two rounds of bead clean-up (AMPure Beads) and assessed resulting gDNA quality using Nanodrop2000 and Qubit 2.0 Fluorometer (dsDNA HS assay) to obtain a minimum ratio of 0.6.

For Oxford Nanopore Technologies (ONT) Nanopore sequencing, we extracted HMW gDNA using a magnetic bead-based protocol described in [86]. We subsequently size selected the gDNA for fragments ≥ 40 kb using a PippinHT (Sage Science).

#### PacBio HiFi sequencing

We sent the final HMW gDNA sample of ∼100 μL, 451.7 ng/μL in 10 mM TrisHCl (∼45 μg HMW) to the Australian Genome Research Facility Ltd (AGRF), St Lucia, Queensland for HiFi 10-15 kb fragment gDNA Pippin Prep size selection, library preparation and PacBio Sequel II sequencing (SMRT Cell 8M).

#### Hi-C proximity-ligation sequencing

Hi-C library preparation and sequencing was conducted at the Ramaciotti Centre for Genomics using the Phase Genomics Plant kit v3.0. A pilot run on an Illumina iSeq 100 with 2 x 150 bp paired end sequencing run was performed for QC using hic_qc v1.0 (Phase Genomics, 2019) with i1 300 cycle chemistry. This was followed by sequencing on the Illumina NextSeq 500 with 2 x 150 bp paired-end high output run and NextSeq High Output 300 cycle kit v2.5 chemistry.

#### ONT Sequencing

We prepared a long-read native DNA sequencing library according to ONT protocol Genomic DNA by Ligation (SQK-LSK110). We performed sequencing on an ONT PromethION using a FLO-PRO002 R9.4.1 flow cell, with three wash treatments and reloads to maximise output, according to the manufacturer’s Flow Cell Wash Kit (EXP-WSH004). We basecalled the fast5 reads to fastq with Guppy version 6.1.2 (ONT), inspecting the output and quality with NanoPlot [88].

#### Genome size prediction

We computed HiFi CCS read Kmer frequencies using Jellyfish v2.2.10 [89] and KMC v3.1.1 [90], with k=19 and a maximum kmer frequency of 10,000 (-k19 -ci1 -cs10000). We used the GenomeScope v2.0 webserver [48] to predict genome sizes.

We carried out additional genome size prediction using single-copy read depth analysis by DepthSizer v1.4.0 [47]. We mapped HiFi CCS and ONT reads to each genome assembly analysed using minimap2 v2.22 [91], and calculated BAM depth and coverage statistics with Samtools v1.13 [92]. We used single-copy genes identified as “Complete” by Benchmarking Universal Single Copy Orthologs (BUSCO) for each assembly. We generated genome size plots with the ggstatsplot package [93] in R v4.1.0.

#### Genome assembly and Hi-C scaffolding

We assembled the genome with the hifiasm v0.15.5 [42] package using PacBio HiFi reads and integrating Hi-C reads. We independently scaffolded genome outputs using the Aiden Lab pipelines [43,44] (assembly v0.1; Figure S3A and B). The assignment of scaffolds to either Haplotype A or B was determined by hifiasm arbitrarily as the parent trees were not available to be sequenced. The ONT data were assembles with Flye (v2.9) [94], polished with Hypo (v1.0.3) [95] and scaffolded with Hi-C data (Figure S1C & D). To scaffold the genomes, we ran the Juicer pipeline (v1.6) [96] with default parameters. To ensure that all duplicate mapped reads were removed, we renamed the merged_sort.txt output from Juicer and reformatted and renamed the merged_nodups.txt to replicate the format of the original merged_sort.txt with the script “cat merged_nodups.txt |sort -- parallel=16 -k2,2d -k6,6d > merged_sort.txt”. We reran Juicer using the newly created merged_sort.txt with additional parameter “-S dedup” and used the final output with the 3D-DNA pipeline (v180922)[45] with the following parameters “-m haploid --build-gapped-map --sort-output”. After we manually curated the assemblies locally within the Juicebox visualisation software (v1.11.08 for Windows) [44], we resubmitted the revised assembly file to the 3D-DNA post review pipeline with the parameters “-- build-gapped-map --sort-output” for final assembly and fasta files.

#### Assembly curation, filtering, and polishing

We tidied Hi-C scaffolds with Diploidocus (v0.18.0) [47] dipcycle mode, using the HiFi reads for both long reads and high accuracy (kmer) reads (assembly v0.2) with each haplotype filtered independently. We assigned chromosomes with PAFScaff (v0.4.1) [97], mapping on to the *Eucalyptus grandis* (GCF_000612305.1) chromosomes (assembly v0.3), and visually compared the two haplotypes, using SynBad (v0.8.4) [98] and DepthKopy (v1.1.0) [47] as guides. We identified some scaffolding errors, which we manually corrected (assembly v0.4) before a second round of Diploidocus tidy on each haplotype (assembly v0.5). We used DepthCharge (v0.2.0) [99] was used to assess for misassemblies, with none identified, however we failed to close any assembly gaps using LR_Gapcloser (v20180904).

Next, we mapped the HiFi reads onto the diploid assembly with Minimap2 (v2.22) [91] and partitioned by haplotype. We separated non-chromosome scaffolds into contigs ran a third round of Diploidocus tidy on each haplotype using the appropriate subset of haplotype-mapped HiFi reads (assembly v0.6).

We then polished the tidied diploid genome with HyPo (v1.0.3) [95] using the HiFi reads mapped with Minimap2 (v2.22) [91] for both the long read and high accuracy data (assembly v0.7). Finally, we renamed the chromosomes according to synteny with the *Eucalyptus grandis* genome [6] to produce v1.0 of the *M*. *quinquenervia* genome.

#### Genome completeness, validation, and annotation

To determine genome completeness, we used Benchmarking Universal Single Copy Orthologs (BUSCO) (v5.1.2) [52] using the lineage dataset embryophyta_odb10. Additionally, we estimated genome assembly quality (QV) using *k-mer* analysis of HiFi read data by Merqury v1.0 with k = 21 [51].

We used the homology-based gene prediction program GeMoMa (v1.7.1) [53] to annotate the genome, utilising four reference genomes downloaded from NCBI: *Arabidopsis thaliana* (TAIR10.1, GCA_000001735.2), *Eucalyptus grandis* [6] (GCF_000612305.1), *Syzygium oleosum* (GCF_900635055.1) and *Rhodamnia argentea* (GCF_020921035.1). We predicted Ribosomal RNA (rRNA) genes with Barrnap (v0.9) [100] and transfer RNAs (tRNAs) with tRNAscan-SE (v2.05) [101], implementing Infernal (Infernal, v1.1.2) [102] filtering for eukaryotes using the recommended protocol to form the high-confidence set. To generate a custom repeat library, we used RepeatModeler (v2.0.1) [54] following genome masking using RepeatMasker (v4.1.0) [103], both with default parameters. We generated the annotation table using the buildSummary.pl RepeatMasker script.

#### Synteny to other Myrtaceae

We used Chromsyn [46]to investigate synteny of *M*. *quinquenervia* to five chromosome-level Myrtaceae genomes available on NCBI: *Angophora floribunda* (GCA_014182895.1), *Eucalyptus grandis* [6] (GCF_016545825.1), *Rhodamnia argentea* (GCF_020921035.1), *Psidium guajava* (GCA_016432845.1) and *Syzygium aromaticum* (GCA_024500025.1). We ordered the species according to phylogenetic relationships [104].

### NLR Analysis

#### NLR annotation with FindPlantNLRs

We developed a comprehensive pipeline to annotate predicted NLR genes from an unmasked genome fasta file input, named FindPlantNLRs [41]. The complete described protocol including software version, dependencies, HMMs and additional scripts are available on GitHub [41].

#### Classification of annotated NLRs and identification of integrated domains

To identify all classes of annotated NLRs, we developed a script that sorted and classified the “gene” types. We ran the file outputs from FindPlantNLRs with the NLR classification script [41]. To further identify novel predicted integrated domains in the annotated NLRs, we developed a script to search the data based on PFAM domain identities not classically associated with NLRs [41]. Resulting files were then sorted to identify the predicted NLR genes by classification and integrated domains per phased genome. The formatted lists were then input to the web-based site https://sankeymatic.com/build/ to create flow diagrams [62]. For all analyses downstream of the FindPlantNLRs pipeline, we included only full NLR gene models which was defined as those genes containing both an NBARC domain and an LRR domain.

#### NLR cluster, duplicated gene, and ortholog analysis

Clustering analysis was based on previous analyses in *E. grandis* and *A. thaliana* genomes [59,105]. We defined a cluster as a genomic region containing three or more predicted *NLR* genes, each of which less than 250 kb from a neighbouring *NLR* gene and with less than 8 non-*NLR* genes between each *NLR*.

We followed the *E. grandis* definition of class classification of *NLR* [59]. *CNL*-type clusters were defined by those containing at least one gene with a *CNL* domain, and no *TNL* type domains. *TNL-type* clusters were defined as those containing at least one gene with a *TNL* domain, and no *CNL* domains. *NL* clusters were defined by those containing only genes with no N-terminal domains. Mixed type clusters were defined as those containing at least two genes with differing N-terminal domains, or lack of N- terminal domain. We visualised the positions of individual *NLRs* and *NLR* clusters on *M. quinquenervia* chromosomes with ChromoMap [106] using base pair start and end positions.

We investigated genome-wide copy numbers using DepthKopy (v1.1.0) [47] for the HiFi and ONT assemblies, with analysis of the HiFi and ONT read data, examining the BUSCO genes, *NLR* annotations, *NBARC* regions, scaffolds and 100 kb windows across the genome.

We identified ortholog within and between both *M. quinquenervia* haplotypes using Blastall (v2.2.26)[107] using a minimum evalue of 1e-10, followed by filtering out hits which have less than 70% identity and score lower than 900. We also identified ortholog using Orthofinder (v2.4.0) [108] using default parameters and inferring maximum likelihood trees from multiple sequence alignments.

#### Phylogenetic analysis of Melaleuca quinquenervia NLRs

To investigate relatedness among NLR genes, we extracted all NBARC domains from the annotated amino acid files for both sets of scaffolds using the chromosome locations with bedtools (v2.29.2) [109] We included an outgroup of amino acid NBARC domains taken from a subset of functionally validated plant NLRs [32]. We reduced the outgroup set to include NBARC domains from eudicotyledons only and incorporated six CNL, two RPW8 and seven TNL-type NBARC domains. We removed 81 predicted transcripts annotated as t2, retaining only t1 predicted reads, from the phased *M. quinquenervia* data and combined the remaining NLR NBARC domains with the outgroups. We aligned the combined sequences with clustal-omega (v.1.2.4)[110], and inferred the phylogenetic tree with IQ-TREE [111] using the following parameters, -bb 1000 -st AA -m LG. We visualised the resulting newick file with iTOL [112] and colour coded according to NLR clade.

#### Transcript evidence for annotated NLRs in Melaleuca quinquenervia

To test for expression evidence for our annotated NLR genes, we downloaded RNASeq data (NCBI PRJNA357284) from a previous *M. quinquenervia* study that investigated responses to the plant pathogen causing myrtle rust [63]. We mapped all the available RNASeq data to the NLR coding sequences for each haploid genome independently using Hisat2 (v2.1.0) [64] with the parameters “hisat2 -p 16 --summary-file MqA/MqB --trim5 15 --trim3 10 --no-unal -p 16 -S <file.sam>”. We processed the sam file outputs with samtools (v1.9) [92] for sorted and indexed bam files and obtained mapping statistics with samtools idxstats. Finally, we calculated the transcripts per million (TPM) for all predicted NLR genes.

## Author contributions

Planned project: AMM, JGB, PAT, RJE, SHC

Wrote paper: AMM, JGB, PAT, RJE, SHC, BS, AJ

Sampling: AMM, JGB, PAT, SHC

DNA extraction: AMM, PAT, SHC, AJ

ONT sequencing: AJ

Primary genome assembly and annotation: SHC, JGB

Hi-C scaffolding: PAT, SHC

Additional assembly curation and QC: RJE, SHC

Synteny and copy number analysis: RJE

NLR pipeline conceptualisation and development: PAT, BS, ZL, TT

NLR analyses: AMM, PAT

Orthology analysis: AMM, ZL

## Acknowledgements

We thank Matt Coyne, David Laughlin and Scott Jones at the Royal Botanic Garden Sydney who assisted with sampling.

